# Bridging the Genotype and the Phenotype: Towards An Epigenetic Landscape Approach to Evolutionary Systems Biology

**DOI:** 10.1101/004218

**Authors:** J. Davila-Velderrain, E. R. Alvarez-Buylla

**Affiliations:** Instituto de Ecología, Universidad Nacional Autónoma de México, Cd. Universitaria, México, D.F. 04510, México; Centro de Ciencias de la Complejidad (C3), Universidad Nacional Autónoma de México, Cd. Universitaria, México, D.F. 04510, México

## Abstract

Understanding the mapping of genotypes into phenotypes is a central challenge of current biological research. Such mapping, conceptually represents a developmental mechanism through which phenotypic variation can be generated. Given the nongenetic character of developmental dynamics, phenotypic variation to a great extent has been neglected in the study of evolution. What is the relevance of considering this generative process in the study of evolution? How can we study its evolutionary consequences? Despite an historical systematic bias towards linear causation schemes in biology; in the post-genomic era, a systems-view to biology based on nonlinear (network) thinking is increasingly being adopted. Within this view, evolutionary dynamics can be studied using simple dynamical models of gene regulatory networks (GRNs). Through the study of GRN dynamics, genotypes and phenotypes can be unambiguously defined. The orchestrating role of GRNs constitutes an operational *non-linear* genotype-phenotype map. Further extension of these GRN models in order to explore and characterize an associated Epigenetic Landscape enables the study of the evolutionary consequences of both genetic and non-genetic sources of phenotypic variation within the same coherent theoretical framework. The merging of conceptually clear theories, computational/mathematical tools, and molecular/genomic data into coherent frameworks could be the basis for a transformation of biological research from mainly a descriptive exercise into a truly mechanistic, explanatory endeavor.

## Introduction

The mechanistic understanding of the mapping of genotypes into phenotypes is at the core of modern biological research. During the lifetime of an individual, a developmental process unfolds, and the observed phenotypic characteristics are consequently established. As an example, a given individual may or may not develop a disease. Can we explain the observed outcome exclusively in terms of genetic differences and an unidirectional, linear relationship between genotype and phenotype? Researchers in biology have mostly assumed so. Over the last decades, scientists under the guidance of such genetic-causal assumption have struggled with inconsistent, empirical observations. The biological relevance of the phenotypic variability produced during the developmental process itself, and not as the consequence of genetic mutations, has only recently started to be acknowledged [1–5].

Understanding the unfolding of the individuals phenotype is the ultimate goal of developmental biology. Evolutionary biology, on the other hand, is largely concerned with the heritable phenotypic variation within populations and its change during long time periods, as well as the eventual emergence of new species. Historically, population-level models seek to characterize the distribution of genotypic variants over a population, considering that genetic change is a direct indicator of phenotypic variation. Certain assumptions are implicit to such reasoning. Are those assumptions justifiable in light of the now available molecular data and the recently uncovered molecular regulatory mechanisms? What is the relevance of considering the generative developmental sources of phenotypic variation in the study of evolution? The aim of this paper is to highlight how a systems view to biology is starting to give insights into these fundamental questions. The overall conclusion is clear: an unilateral *gene-centric* approach is not enough. Evolution and development should be integrated through experimentally supported mechanistic dynamical models [6–13].

In the sections that follow, we first present a brief historical overview of evolutionary biology and the roots of a systematic bias towards linear causation schemes in biology. Then, we discuss the assumptions implicit in the so-called neo-Darwinian Synthesis of Evolutionary Biology – the conventional view of evolution. In the last section, we briefly describe an emerging research program which aims to go beyond the conventional theory of evolution, focusing on a nonlinear mapping from genotype to phenotype through the restrictions imposed by the interactions in gene regulatory networks (GRNs) and its associated epigenetic landscape (EL). Overall, this contribution attempts to outline how the orchestrating role of GRNs during developmental dynamics imposes restrictions and enables generative properties that shape phenotypic variation.

## Darwin’s Legacy

Darwin eliminated the need for supernatural explanations for the origin and adaptations of organisms when he put evolution firmly on natural grounds [14]. In the mid-19th century, Darwin published his theory of natural selection [15]. He proposed a natural process, the gradual accumulation of variations sorted out by natural selection, as an explanation for the shaping and diversity of organisms. This insight was what put the study of evolution within the realms of science in the first place [14]. Although it has had its ups and downs [16], the Darwinian research tradition predominates in modern evolutionary biology. Much of its success is due to a new (gene-centric) interpretation, the so-called neo-Darwinian modern synthesis [17]: the merging of mendelian genetics and Darwin’s theory of natural selection due to prominent early 20th century statisticians. In this framework, development was left outside, and evolution is seen as a change in the genotypic constitution of a population over time. Genes map directly into phenotypes (see Figure 1a), implicitly assuming that genetic mutation is the prime cause of phenotypic variation. Observed traits are generally assumed to be the result of adaptation, the process whereby differential fitness (the product of the probability of reproduction and survival) due to genetic variation in a particular environment, leads to individuals better able to live in such an environment.

**Figure 1.**
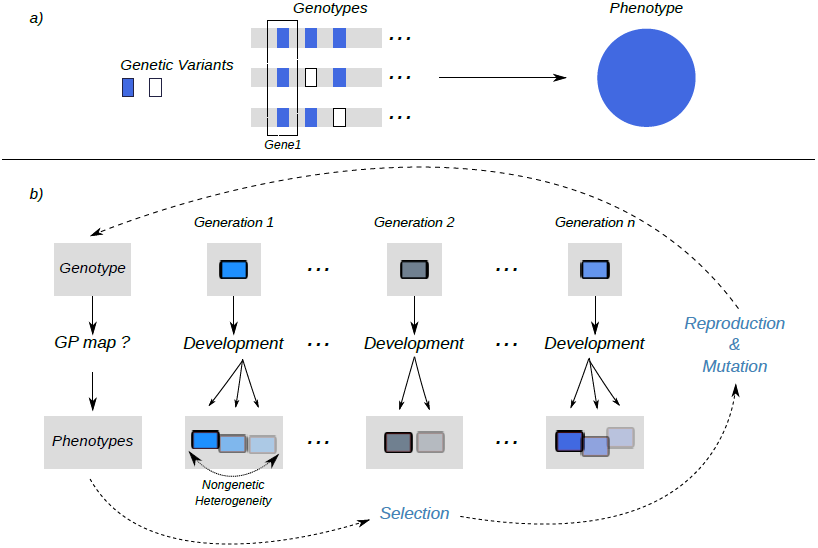
a) A straightforward genotype-phenotype relationship: the genetic distribution of the observed locus would completely mirror the phenotypic distribution; gene interactions are ignored; as a result, three different genotypes would correspond to the same phenotype given the locus under observation. b) A developmental process from genotype to phenotype, a GP map: through the development of an individual nongenetic phenotypic variation is generated each generation; in an evolutionary time-scale, evolution operations (blue) produce genetic variation. Selection acts on phenotypes; phenotypic variation is the product of both genetic mutational operations and epigenetic developmental processes.

### From Natural Selection to Natural Variation

Natural selection - a force emanating from outside the organism itself - is the conceptual core of the Darwinian research tradition. Conceptually, the general process is as follows. *Random* mutations occur during reproduction; these mutations are responsible for generating different (genetic) types of individuals. The selection process then results from the fact that each type has certain survival probability and/or is able to achieve certain reproductive performance given the environment. Through this differential rate, some types are maintained while others are dismissed. It is said that, in this way, selection makes a “choice” [18]. From a wider perspective, it is generally accepted that selection is a generic process not restricted to biological evolution [19]. Any error-prone communication process in which information is consequently transmitted at different rates leads itself to a selection mechanism. However, despite the appealing conceptual clarity of the selection mechanism, it is not generally appreciated that the complexity inherent to biological systems hinders the mechanistic understanding of biological evolution. Because the reproductive performance of a given type of variant is, mainly, a function of its phenotype; the paradigmatic selection process described above is plausible when one assumes a straightforward causation of phenotype by genotype [10]. A more faithful model of biological evolution should explicitly consider a genotype-phenotype (GP) map [20,21], a developmental mechanism which specifies how phenotypic variation is generated (Figure 1b). The generated variation is then what triggers selection [22]. Importantly, a deviation from a linear causation view of development would potentially impact the rate and direction of evolution [8,23,24].

Although not always discussed, Darwin himself devoted much more attention to variation than to natural selection, presumably because he knew that a satisfactory theory of evolutionary change requires the elucidation of the causes and properties of variation [25]. After all, natural selection would be meaningless without variation. Ironically, given the success of the neo-Darwinian framework, phenotypic variation to a great extent has been neglected in the study of evolution [26]. The mechanistic understanding of the sources of phenotypic variation constitutes a fundamental gap in conventional evolutionary theory. Neither Darwin, nor the founders of the neo-Darwinian modern synthesis were able to address this problem given the biological knowledge available at the time. Moreover, deviations from the basic assumptions of the conventional theory were not always generally appreciated [27].

### Implicit Assumptions in Evolution

Being the development of science an evolutionary process itself, it is reasonable to expect that social-historical contingency has profoundly biased the pathways of scientific inquiry. This seems to be the case in the history of biology. For example, (1) Darwin’s war against divine explanations for biological complexity caused within the scientific community an automatic rejection for any goal-oriented activity within organisms. This situation favored the adoption of the the idea of random (uniform) variation [28,29]. (2) The mainstream focus of neo-Dawinism on optimizing reproductive success (fitness) by natural selection of random variants; on the other hand, implicitly neglected the relevance of gene interactions (see Figure 1a) [30]. Finally, (3) the establishment of the central dogma of molecular biology (gene → mRNA → protein) further cemented a linear, unidirectional scheme of causation of molecular traits (one gene - one protein, one trait) [10]. These events are thought to be associated with a deeply rooted systematic bias towards linear causation schemes in biology [10,31]. They also favored the adoption of three major implicit assumptions upon which the neo-Darwinian tradition was developed, namely: (1) mutational events occur randomly (e.g. unstructured) along the genome; (2) given that the phenotypic effects of successive mutations in evolution are of additive nature, gene interactions and their phenotypic influence can be, to a large extent, ignored; and (3) the phenotypic distribution of mutational effects mirrors the genetic distribution of mutations [30].

Scientists are now re-examining the most basic assumptions about evolution in light of post-genomic, systems biology [28,32]. Compelling evidence has been presented even against assumption (1) above. For example, Shapiro has shown how a truly random (unstructured) nature of mutational events is empirically unsustainable. He has coined the term “natural genetic engineering”, referring to the known operators that produce genomic changes and which are subjected to cellular regulatory regimes of epigenetic character [29]. It seems that the generative properties of genetic variation are nonuniform, and thus, biased as well. Assumptions (2) and (3) above are, instead, mainly concerned with how phenotypic variation is generated given a genetic background; or in other words, with the mechanistic understanding of the GP map. Here, we are concerned with this developmental process and its evolutionary relevance.

## From Genes to Networks

At the beginning of the 21th century, biology confronted an uncomfortable fact: despite the increasing availability of whole-genome sequence data, it was not possible to predict, or even clarify, phenotypic observations. In fact, we now know that there is not sufficient information in the linear DNA sequences of the complete genomes to recover and/or understand the diverse phenotypic states of an organism. It was clear that cell behavior was much more complex than anticipated. Since then, biological research has increasingly been oriented towards a systems-level approach that goes beyond obtaining and describing large data sets at the genomic, transcriptomic, proteomic or metabolomic levels. An assumption of such *systems* approach to biology is that cell behavior can be understood in terms of the dynamical properties of the involved molecular regulatory networks. Modern molecular evolutionary studies are starting to incorporate this network thinking: genes are not individual entities upon which evolutionary forces act independently. Evolutionary forces, functional constraints, and molecular interactions are conditionally dependent on the systems level [33]. How a systems-view impacts our understanding of the GP map?

### Fundamental Sources of Natural Variation

Although the concepts of genotype and phenotype are fundamental to evolution, it is not straightforward to operationally define them: In practice genotype and phenotype distinctions are just partial [34]. This is partially the reason why simple theoretical models are so important for the epistemology of evolution. A common working model in systems biology is that in which the phenotypic state is defined at the cellular level. The cellular phenotype is represented by the activity of each of its genes, its expression pattern. Since the regulatory interactions among the genes within the cell constitute a network, the network effectively represents the genotype of the cell, while its associated expression profile represents its phenotype (Figure 2). The structure of the former derives directly from the genome, while the latter changes through development. In practice, we just observe certain expression patterns (e.g cell-types) - with small deviations - and not others. Why is that?

**Figure 2.**
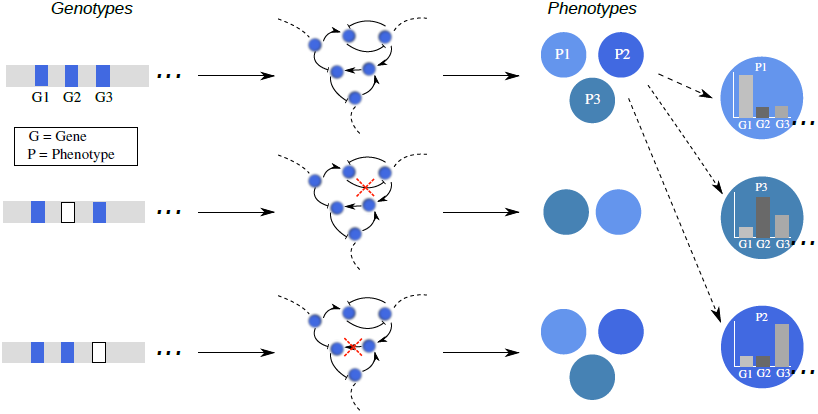
The orchestrating role of GRNs constitutes a *non-linear* GP map. Through the restrictions imposed by the interactions in GRNs, cellular phenotypes (represented by expression profiles) are generated. Due to the nonlinear character of GRN dynamics, the GP map is one-to-many. The effect of mutations in the phenotype is not uniformly distributed over the genes, but depends on the interactions: mutations can or cannot result in different phenotypes depending on the genetic background and the location of the affected genes in the network.

#### GRN developmental dynamics generates phenotypic nongenetic (epigenetic) heterogeneity

When thinking in terms of a genotype-phenotype distinction based on GRN dynamics, it is natural to consider an abstract space where all the virtually possible phenotypes reside. We call this space the *state-space*. Empirical observations suggest that something should be maintaining cells within specific, restricted regions of this space. The structured nature of the underlying GRN determines a trajectory in this state-space: given the state of the genes regulating a gene *i*, and the functional form of the regulation, the gene *i* is canalized to take specific future states. Eventually, this self-organizing process would inevitably lead to the establishment of those states which are logically consistent with the underlying regulatory logic. In this way, the GRN imposes constraints to the behavior of the cell. The resultant states are denominated *attractors* and correspond to observable cell-types. These are the basis of the well developed dynamical-systems theory of cell biology (for a review, see [35,36]). This theory was first applied to propose a GRN grounded on experimental data for understanding how cell-fate specification occurs during early flower development (see, [37,38] and update in [39]). Originally, the approach was inspired by theoretical work in randomly assembled networks by Stuart Kaufman [40]. In the last decades, the theory has been supported by a wealth of consolidated theoretical and experimental work (see, for example [7,13,41]).

Through GRN dynamics, development generates cellular phenotypes. The general acceptance of this generative role necessarily implies deviations from the neo-Dawinian framework. Importantly, (1) the effect of a perturbation (mutational or otherwise) on the manifested phenotype is not uniformly distributed (truly random) across all the genes in the network, and (2) the interactions in the network are fundamental to the establishment of the phenotype. The orchestrating role of GRNs constitutes a *non-linear* GP map: phenotypic variation does not scale proportionally to genotypic variation; it is not linear (Figure 2). Two important consequences of these mechanistic view of developmental dynamics have been eloquently pointed out recently. First, the nonlinear character of this mapping ensures that the exact same genotype (network) is able to produce several phenotypes (attractors) [40]. Second, given that molecular regulatory events are stochastic in nature, a cell is able to explore the state-space by both attracting and dispersing forces - forces that slightly deviate the dynamics from the determined trajectory. Any phenotype of a cellular population at any given time is statistically distributed [10]. These sources of variation are the natural product of developmental dynamics. Consequently, at any given time, a population can manifest phenotypic variation that is relevant to evolution (heritable) in the absence of genetic variation. How can we study evolution without ignoring the fundamental role of developmental dynamics?

## Evolutionary Systems Biology Approaches

A systems view to evolutionary biology, in which network models as GP mappings are considered explicitly, is under development (see, for example [9,11,42]). Within this general framework, several specific approaches are proposed in order to study the evolutionary consequences of considering developmental sources of phenotypic variation. In this section, we briefly present a preview of an emerging complementary approach.

### Epigenetic(Attractors) Landscape Evolution

In 1950s, C.H. Waddington proposed the conceptual model of the epigenetic landscape (EL), a visionary attempt to synthesize a framework that would enable an intuitive discussion about the relationship between genetics, development, and evolution [43]. His reasoning was based on the consideration of a fact: the physical realization of the information coded in the genes - and their interactions - imposes developmental constraints while forming an organism. Now, in the post-genomic era, a formal basis for this metaphorical EL is being developed in the context of GRNs [10,44,45]. The key for this formalization is an emergent ordered structure embedded in the state-space, the attractors landscape (AL). As well as generating the cellular phenotypic sates (attractors), the GRN dynamics also partitions the whole state-space in specific regions and restricts the trajectories from one state to another one. Each region groups the cellular states that would eventually end up in a single, specific attractor. These sub-spaces are denominated the attractor’s *basin* of attraction. Given this (second) generative property of GRN dynamics, the formalization of the EL in this context is conceptually straightforward: the number, depth, width, and relative position of these basins would correspond to the hills and valleys of the metaphorical EL. We refer to this structured order of the basins in state-space as the AL (see Figure 3). The characterization of an AL would correspond, in practical terms, to the characterization of an EL. Is this formalized EL useful for the mechanistic understanding of phenotype generation?

**Figure 3.**
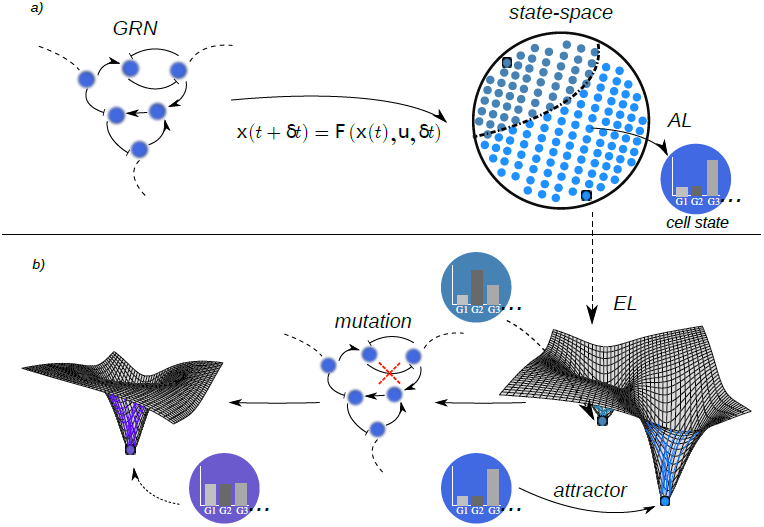
The Epigenetic (Attractors) Landscape. a) Through a dynamical mapping - a mathematical representation of the gene regulatory logic - GRNs generate both the cellular phenotypes (attractors) and the ordered structure of the state space - the AL. Through the structure of the AL, the EL is formalized in the context of GRNs. b) The number, depth, width, and relative position of attractors correspond to the hills and valleys of the EL. The topography of the landscape can change in response to perturbations. Mutations could eventually reshape the EL and consequently eliminate and/or generate novel phenotypes.

### Multicellular morphogenetic processes unfold naturally in the EL

The structured EL is a generative property of the GRN dynamics, but at the same time, it also constrains the behavior of a developing system. While a developing system is following its dynamically constrained trajectory in state-space, developmental perturbations from internal or external origin can deviate it. In a cellular population, then, the probability of one phenotypic transition or another during development, as well as the stationary distribution of phenotypes, would be conditioned on both the localization of the individual cells in the EL and on the landscape’s structure. As a general result of this interplay, determinism and stochasticity are reconciled, and robust morphogenetic patterns can be established by a hierarchy of cellular phenotypic transitions (see, for example [44,45]). In this way, morphogenetic processes effectively unfold on ELs. How could this theoretical framework improve the understanding of evolutionary dynamics?

We have an effective nonlinear GP map from GRN to EL. Given an experimentally characterized GRN, the EL associated to real, specific developmental processes can be analyzed ([13,44,45]). Both cellular phenotypes (attractors) and morphogenetic patterns are linked to the structure of the EL. Can we describe this structure quantitatively? How robust is the structure to genetic (network) mutation? Can we describe quantitatively the change in structure in response to both mutational and developmental perturbations? How slower is this rate of change in comparison to the time-scale of developmental dynamics (landscape explorations)? What are the phenotypic consequences of different relative rates of change? Does the resultant evolutionary trajectory of the reshaped EL structure subjected to mutations predicts the probability of phenotypic change (innovation) - based, for example, in the appearance of new cellular phenotypes or morphogenetic patterns? (Figure 3). Insight into these and similar questions could enhance the mechanistic understanding of the evolution of morphogenetic processes.

## Conclusion and Challenges

A modern systems view to biology enables tackling foundational questions in evolutionary biology from new angles and with unprecedented molecular empirical support. Little is known about the mechanistic sources of phenotypic variation and its impact to evolutionary dynamics. The explicit consideration of these processes in evolutionary models directly impacts our thinking about evolution. Simple, generic dynamical models of GRNs, where genotypes and phenotypes can be unambiguously defined, are well-suited to rigorously explore the problem. Further extension of these models in order to explore and characterize the associated EL enables the study of the evolutionary consequences of both genetic and non-genetic sources of phenotypic variation within the same coherent theoretical framework. The network-EL approach to evolutionary dynamics is promising, as it directly manifests the multipotency associated with a given genotype. Although conceptually clear and well-founded, its practical implementation implies several difficulties, nonetheless; specially in the case of high dimensional systems. Work has been done in which the landscape associated with a specific, experimentally characterized GRN is described quantitatively in terms of robustness and state transition rates [46], for example. However, neither the methodology to derive ELs from GRNs, nor the quantitative description of ELs are standard procedures. Most approaches require approximations and are technically challenging for the case of networks with more than 2 nodes. Further research in the quantitative description of experimentally grounded GRNs is still needed in order to explore the constraints and the plasticity of ELs associated with a genotypic (network) space. In this regard, discrete dynamical models are promising tools for the exhaustive characterization of the EL, and for the study of multicellular development [45]. A second major challenge is the generalization of GRN dynamical models in order to include additional sources of constraint during development. Tissue-level patterning mechanisms such as cell-cell interactions; chemical signaling; cellular growth, proliferation, and senescence; inevitably impose physical limitations in terms of mechanical forces which in turn affect cellular behavior. Although some progress has been presented in this direction [47,48], the problem certainly remains open.

The post-genomic era of biology is starting to show that old metaphors such as Waddington’s EL are not just frameworks for the conceptual discussion of complex problems. The merging of conceptually clear theories, computational/mathematical tools, and molecular/genomic data into coherent frameworks could be the basis for a much needed transformation of biological research from mainly a descriptive exercise into a truly mechanistic, explanatory and predictive endeavor - EL models associated with GRNs being a salient example.

